# Perspectives in conducting task-based research in pediatric surgical epilepsy patients

**DOI:** 10.64898/2026.07.02.734030

**Authors:** Jacob Leisawitz, Sandra F. Georges, Alyssa M. Field, Saman Asghar, Gabrielle Foox, Andrew J. Watrous, Howard L. Weiner, Anne E. Anderson, Liberty S. Hamilton

## Abstract

**Objective:** Pediatric epilepsy patients undergoing stereo-electroencephalography (sEEG) for ictal onset evaluation provide a rare window to study the developing brain. While methodological frameworks for task-based sEEG research are well-established in adults, pediatric-specific guidance remains underdeveloped. Furthermore, many pediatric epilepsy patients have comorbidities that might typically exclude them from participating in research. We examine factors that influence research participation and discuss considerations for conducting sEEG research in children.

**Methods:** Here, we present a retrospective analysis of task-based research participation patterns from an NIH-funded study of speech and language representations (1R01DC018579) in 66 patients (ages 4-24) undergoing sEEG monitoring at Texas Children’s Hospital to determine whether specific comorbidities influenced research participation.

**Results:** Eighty-nine percent (n=66) of patients approached for consent agreed to participate in the study. Despite high rates of comorbidities including neurocognitive disorder (66.67%), language delay (31.75%), global developmental delay (23.81%), mood disorders (33.33%), ADHD (46.03%), autism spectrum disorder (14.29%) or other cognitive/intellectual disabilities (36.51%), all participants engaged in at least one task. While the majority of these diagnoses did not appear to influence subject participation, global developmental delay was associated with a significant reduction in time spent on active tasks.

**Discussion:** Despite high prevalence of neuropsychological comorbidities among participants, our evidence suggests that these participants contribute meaningfully to studies investigating important developmental questions. We suggest strategies for tailoring task-based research to accommodate the unique needs of individuals in this population. Such practices are important for ensuring that research studies reflect the true diversity of the population.

## 1. Introduction

### 1.1 Epilepsy and stereo-electroencephalography (sEEG)

Approximately 0.6% of children aged 0-17 years have active epilepsy [1,2]. Between 15 and 25% of these children exhibit drug-resistant epilepsy (DRE) [3]. Due to uncontrolled seizures, effects on development, and the impact on quality of life, these patients are often referred for epilepsy surgery [4–8]. This typically involves a dual-phase workup. Phase 1 utilizes non-invasive tests to inform planning for Phase 2 based on predicted epileptogenic zones (EZ) [9]. Phase 2 can involve surgical placement of intracranial electrodes within potential EZs, known as stereo-electroencephalography (sEEG) [9]. Following sEEG electrode implantation, patients are monitored in the Epilepsy Monitoring Unit (EMU) for 1-4 weeks to localize and define the seizure onset zone (SOZ).

sEEG recordings performed as part of epilepsy surgery offer a unique opportunity to study the brain at a high spatiotemporal resolution [10,11]. Such recordings in adults have been used to understand neural processing of speech [12–20], attention [21–25], vision [26], memory and spatial navigation [27–30], and for building brain computer interfaces [31–36]. In pediatric participants, these recordings enable researchers to study the developmental trajectory of brain function [37–41]. However, many comorbidities and cognitive difficulties exhibited by pediatric patients with DRE are exclusion criteria for research studies. In addition, the nature of this research and the vulnerability of the participant population require sensitivity by the research team.

### 1.2 Comorbidities in Pediatric Epilepsy Patients

Common comorbidities in children with DRE include attention deficit hyperactive disorder (ADHD), autism spectrum disorder (ASD), developmental and language delays, intellectual disability, mood disorders such as anxiety and depression, and various other neurocognitive disorders [42–44]. This poses unique challenges to researchers for consenting, task-design, and maintaining participant engagement.

### 1.3 Guidelines and Considerations for Intracranial Research

Guidelines for intracranial research have been discussed by various groups [45–49]. Recurring themes include 1) prioritization of clinical care, 2) informed consent procedures that allow for patient autonomy and continued consent, and 3) establishing the role of the researcher as separate and distinct from healthcare providers. Still, pediatric-specific guidelines are underdeveloped. Here, we present our perspectives on implementing task-based intracranial research studies in children. We offer recommendations for creating inclusive, developmentally appropriate research paradigms that effectively engage a diverse group of pediatric participants. Finally, we analyze the impact of pediatric comorbidities on research participation.

## 2. Methods

### 2.1 Parent Study

#### 2.1.1 Structure

We conducted a retrospective secondary analysis of data collected during an NIH-funded investigation of the neural mechanisms underlying speech processing in the developing brain.

The parent study (NIH grant R01DC018579) was performed at Texas Children’s Hospital (TCH) in Houston, Texas. Participants engaged in tasks designed to probe speech perception, speech production, and auditory or visual attention. Descriptions of each task are provided in Supplementary Materials Appendix A.

#### 2.1.2 Participants

74 patients and their families were approached for participation in the TCH Epilepsy Monitoring Unit (EMU) for Phase 2 evaluation. 66 (89%) consented to participate, including three patients during two separate admissions, yielding 63 unique participants. Eight patients declined participation; reasons for declining consent were not reported. Demographic characteristics of patients as well as comorbidities are detailed in **Table 1**. Inclusion criteria were: (1) age 4–24 years, (2) pre-operative diagnosis of intractable epilepsy, (3) admission for intracranial EEG monitoring, and (4) primary language of English. All procedures were approved by the University of Texas at Austin Institutional Review Board under a single-IRB agreement with Baylor College of Medicine. For patients <18 years of age, parental permission and age-appropriate assent was obtained. For patients >18, written informed consent was obtained. On average, consent was obtained on post-operative day (POD) 2.6 and data collection began on POD 3.8. This gave patients and families time to recover from the immediate effects of surgery before deciding whether to participate in research.

**Table 1.** Participant characteristics. ADHD = attention deficit hyperactivity disorder, ASD = autism spectrum disorder, GDD = global developmental delay, LD = language delay, MD = mood disorder (anxiety and/or depression), NCD = mild, moderate, or major neurocognitive disorder. “Other” included 1) Intellectual Disability (n=8), 2) Specific Learning Disorder (n=6), 3) Executive Function Deficit (n=5), 4) Borderline Intellectual Function (n=3), 5) Attention and Concentration Deficit (n=2), 6) Global Dysfunction (n=2). All diagnoses were made by a board-certified pediatric clinical neuropsychologist, with relevant ICD-10-CM codes given where available. *The developmental stage did not change between admissions for any of the 3 patients enrolled twice.

| Characteristic | n out of 63 (%) |
| --- | --- |
| Age/Developmental Stage* | Early Childhood (Age 2-5): 5 (8.0%) |
|  | Middle Childhood (Age 6-11): 23 (36.5%) |
|  | Early Adolescence (Age 12-17): 23 (36.5%) |
|  | Late Adolescence (Age 18-24): 12 (19.0%) |
| Sex | Male: 32 (50.8%) |
|  | Female: 31 (49.2%) |
| Race/Ethnicity | Asian, Not Hispanic or Latino: 3 (5%) |
|  | Black, Not Hispanic or Latino: 4 (6%) |
|  | White, Hispanic or Latino: 21 (33%) |
|  | White, Not Hispanic or Latino: 32 (51%) |
|  | More than one race, Not Hispanic or Latino: 3 (5%) |
| Comorbid Diagnoses | NCD: 42 (66.6%) – ICD codes G31.84, F02.81, F71 |
|  | ADHD: 29 (46.0%) – ICD codes R41.840, F90.0, F90.2, F90.9 |
|  | MD: 21 (33.3%) – ICD codes F43.23, F41.9, F43.23, F33.41, R45.89, F32.9, F32.1 |
|  | LD: 20 (31.7%) – ICD code F80.2 |
|  | GDD: 15 (23.8%) – ICD code R62.50, F88, R62.50 |
|  | ASD: 9 (14.3%) – ICD code F84.0 |
|  | Other: 23 (36.5%) |
|  | None: 3 (4.8%) |

#### 2.1.3 Task Structure and Modifications

Prior to approaching patients, research staff reviewed neuropsychology reports to tailor experimental task priority to the patient’s developmental and cognitive profile. Tasks were categorized as passive or active: passive tasks only required participants to listen to auditory stimuli and/or look at a screen, while active tasks required subjects to 1) listen to and repeat words, phrases, and sentences, or 2) follow on-screen directions to play a game that included button presses. Short data collection sessions (e.g. <1 hour) were conducted across multiple days and scheduled around clinical care and patient comfort. If seizures occurred, sessions were stopped or rescheduled. When possible, participants were allowed to choose the order and duration of tasks to match their comfort and energy level. Tasks were designed to be developmentally appropriate and engaging for a wide age range and for participants with varying cognitive and behavioral abilities. Specific modifications for achieving this are detailed in the Discussion.

### 2.2 Chart Review

Consented participants’ age, sex, bilingual status, time spent completing research tasks, and history of neuropsychological diagnoses were compiled. All patients underwent neuropsychological testing as part of their initial workup before undergoing epilepsy surgery. Therefore, diagnoses were made by a licensed clinical neuropsychologist (example ICD codes are given in Table 1). Diagnoses considered included neurocognitive disorder (NCD), attention-deficit-hyperactivity disorder (ADHD), mood disorder (MD), language delay (LD), global developmental delay (GDD), and autism spectrum disorder (ASD).

### 2.3 Statistical Analysis

We examined effects of individual comorbidities on task participation. Because data were not normally distributed, we used a nonparametric Mann-Whitney unpaired t-test to identify statistically significant differences in participation time between comorbidity groups. Statistical tests were performed using GraphPad Prism (v10.6.1). We fit a linear mixed effects model using the lmerTest library [50] in RStudio (v2023.06.0+421, Posit Software) to test whether task type (passive vs. active) and number of comorbidities affected time spent in the task, with participant as a random effect.

## 3. Results

### 3.1 Practical and Ethical Considerations for Intracranial Research in Children

#### 3.1.1 Consent, Continued Consent, and Data Collection

Maintaining participant agency is an essential aspect of ethical human subjects research. Because patients are often weaned off AEDs and sleep-deprived to induce seizures, their medical condition can vary from day to day. Researchers should thus gauge the willingness of the patient and caregivers to participate before each session, reiterating that participation is voluntary, and attending to body language, mood, and engagement. Caregivers, who may be working remotely or hosting visitors, should be consulted about preferred timing and about any religious and/or social restrictions relevant to task selection.

When possible, we allowed participants and caregivers to choose tasks according to preference. When specific tasks are desired due to electrode coverage, we explained our goals for the session, aligning our research aims with participant engagement. A participant with dense motor cortical coverage, for example, would be a strong candidate for active speaking tasks, but these could be alternated with easier listening tasks to give the patient a rest.

#### 3.1.2 Prioritizing Clinical Care

Successful study participation requires a clear understanding of the boundary between research and clinical care. During the consent process, it is important that the research team clearly communicates that participation in research will not interfere with and is separate from their clinical care [45]. Researchers should be aware of the overall spectrum of the patient’s clinical care, including when the attending physician and team have rounded, plans for cortical stimulation and functional mapping, and the trajectory of their treatment. Researchers should not wake patients when asleep or enter the room when any member of the clinical team is present.

#### 3.1.3 Additional Considerations for Children

Conducting intracranial studies in children poses unique challenges for researchers as compared to the same studies in adults. Children may differ from adults in terms of their motivation to engage, their desire to be in control, and their ability to maintain engagement. Adults can more readily grasp abstract reasons for participating despite having no direct benefit, whereas children, especially younger ages who are cognitively impaired, may require different strategies to motivate engagement. For example, rather than telling a 6-year-old child that we are doing a research study to learn about speech representations in the developing brain, it is often more effective to describe to them the task in a way they understand, and that frames their participation as fun and easy. For example, for introducing our movie trailer watching task for a younger child (see Supplementary Materials Appendix A), it may be effective to say “We’re going to watch some movies on our iPad today! How does that sound? Do you like Moana?”

Another important consideration for researchers is that children may have a stronger need to feel a sense of control, as opposed to an adult who may be able to adapt to less engaging research tasks. A strategy we have found to be useful to allow the subject to feel a sense of autonomy has been utilizing bargaining tactics in which researchers offer task choices rather than ultimatums. Take, for example, a child who only wants to watch “Trolls”, but the researchers have already collected multiple repetitions of this trailer and want to diversify their dataset. It may be useful to propose something like the following, “Today, we are going to take turns picking movies. When it’s your turn, you can pick Trolls, or any other movie you want. Then it will be my turn to pick. You can go first.” A similar framing can introduce a less familiar or more difficult task as a game. Researchers should remember to allocate time in each session for these types of “bargaining” conversations with the participant.

The assistance of a parent or legal guardian in encouraging or reassuring their child can be helpful for researchers. This has helped with initiating data collection on the first day, as it is not uncommon for children to be apprehensive towards researchers at first. In cases where the child is feeling ill, or simply does not want to participate, the researcher should terminate the session.

### 3.2 Role of Comorbidities

#### 3.2.1 Chart Review

Almost all participants (95.2%) had at least one comorbid neuropsychiatric diagnosis (Table 1). Disorders such as ADHD and ASD, known to present significant challenges in maintaining attention and focus, were present in 46% and 14.3% of our study participants, respectively. Mood disorders such as anxiety and depression were diagnosed in 33.33% of consented participants. Global and language delays are also prominent amongst children with DRE and were diagnosed in 23.8% and 31.8% of our study participants, respectively. Finally, neurocognitive disorder (NCD), the most broad and prevalent diagnosis in this research population, was found in 66.7% of our study participants.

#### 3.2.2 Effect of Individual Comorbidities on Subject Participation

We analyzed whether comorbid diagnoses affected overall participation, or the percentage of total research time participants spent completing active tasks. None of the comorbidities examined significantly affected the total duration of patient participation in research activities (**Figure 1**). Only global developmental delay was associated with a difference in the proportion of time spent on active research tasks (p = 0.035).

**Figure 1.**
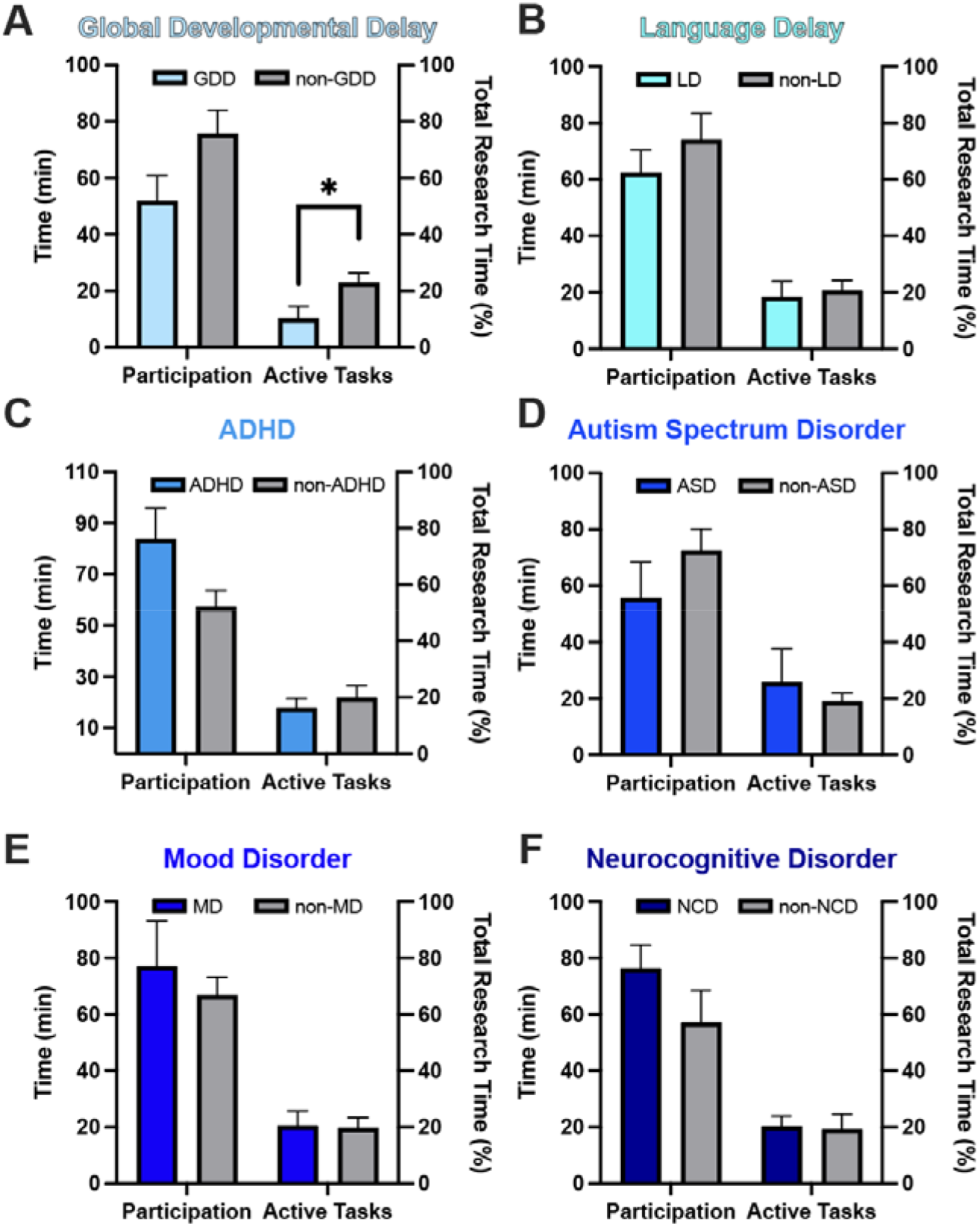
Effect of comorbidities on research participation and percentage of total research time spent doing active tasks.Data are reported as mean + SEM for both total participation time, and percentage of time spent in active tasks.

In our study, patients could participate in several independent tasks, most of which were used for separate manuscripts related to the parent grant. Thus, although patients could theoretically complete all of our tasks, this was not required by our protocol, and instead participants could complete any of the tasks based on their electrode coverage, cognitive ability, and interest. All participants included in this analysis contributed at least one analyzable data block, defined as the successful completion of at least one task (see Table 2). Only one patient discontinued tasks prematurely due to fatigue. Sessions lasted from 5–115 minutes (mean = 28 minutes) and were usually distributed over multiple days, with the average participant completing 2.67 sessions during their admission. Passive listening tasks were most frequently completed by our patients. Despite passive tasks representing 70% of total completed tasks, at least one active task was completed by 65.15% of participants, while 42.42% of participants completed 2 or more active tasks.

**Table 2.** Number of experimental sessions, task completion, and active task engagement across subjects with different comorbidities.

|  | GDD | LD | ADHD | ASD | MD | NCD |
| --- | --- | --- | --- | --- | --- | --- |
| Sessions / Subj. (Mean SEM) | 2.2 0.3 | 2.7 0.3 | 3.1 0.4 | 1.9 0.3 | 2.9 0.6 | 2.8 0.3 |
| Tasks / Subj. (Mean SEM) | 4.9 1.1 | 6.5 1.1 | 7.8 1.1 | 5.9 1.8 | 7.7 1.5 | 7.7 1.0 |
| Session Length (Mins; Mean) | 26.8 2.7 | 26.0 1.7 | 30.7 1.7 | 32.1 4.2 | 28.4 1.6 | 29.9 1.4 |
| Subjs. Completing >1 Active Task (%) | 40 | 59 | 69 | 56 | 73 | 69 |
| Subjs. Completing >2 Active Tasks (%) | 20 | 32 | 53 | 33 | 59 | 44 |
| Length of Stay (Days; Mean) | 12.4 | 9.7 | 11.8 | 9.4 | 11.9 | 11.8 |

#### 3.2.3 Effect of Multiple Comorbidities on Subject Participation

We examined whether time spent performing research tasks depended on number of comorbidities and task type (active vs. passive) (**Figure 2**). Passive tasks had significantly higher participation time than active (p=5.24×10^−9^, df=64, t-value=6.743). There was no effect of number of comorbidities on time spent in either task type (passive or active; p=0.442, df=63, t-value=0.774). By modifying our task designs (see Discussion), we were able to include participants with a broad spectrum of abilities.

**Figure 2.**
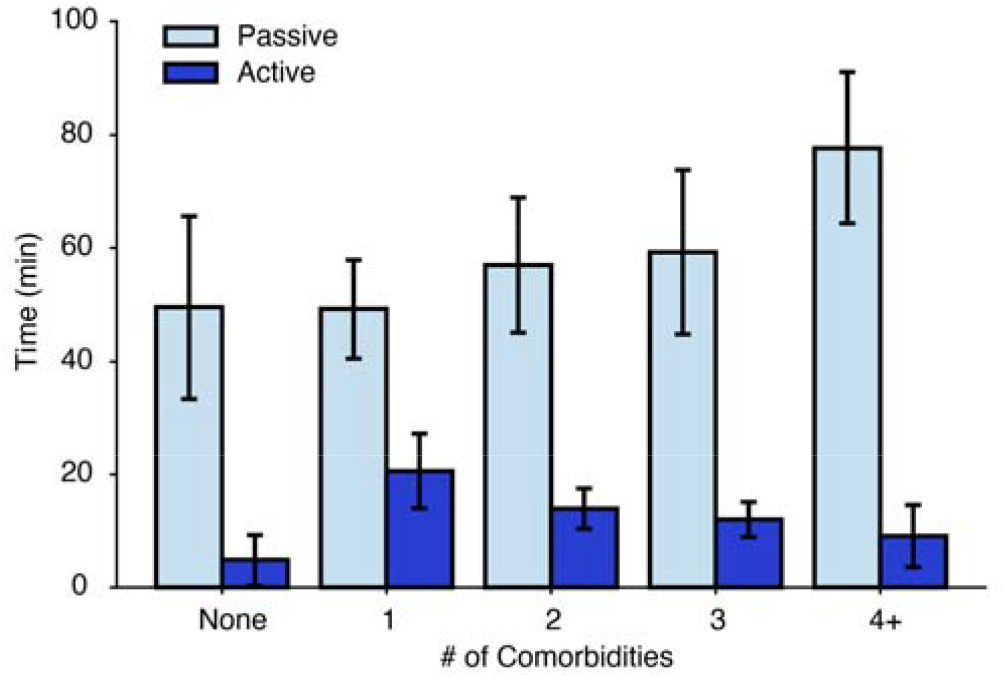
Time spent completing passive and active tasks amongst participants with an increasing number of comorbid diagnoses. Data are expressed as mean + SEM of minutes spent completing tasks in each group

#### 3.2.4 Case Studies

Two case studies are provided below as examples of task modifications and observations from patients with multiple comorbidities.

**Case 1:** A 4-year-old female with a history of language delay and epilepsy of unclear etiology but a suspected genetic cause had sEEG electrodes placed bilaterally in frontal and parietal regions and right-sided temporal lobe. Due to her age, she participated in only passive tasks, which included watching movie trailers in Spanish and English, as well as the “Frog Story” (FS) listening task (see Appendix A). The movie trailer and FS tasks were adopted to replace the TIMIT sentence corpus. Although TIMIT is well-validated scientifically [51], it was poorly tolerated by many participants due to its length and lack of narrative. The FS and movie trailer tasks allowed us to assess receptive language responses in a more accessible and enjoyable format. We have shown in other work that the movies can provide comparable results to TIMIT on acoustic and phonetic selectivity in the brain (Desai et al. 2021). In this case, the patient was very engaged and completed a total of 138 minutes of passive tasks.

**Case 2:** An 11-year-old male with prior right frontal craniotomy and resection underwent sEEG implantation with diffuse right frontal coverage, including supplementary motor areas along the posterior resection margin. His neuropsychological evaluation documented ADHD and relative difficulties with sustained attention and verbal expression. Task selection was informed by his neuropsychological profile and discussions with his mother regarding his interests. His mother shared that he enjoyed computer and video games but was often reserved when asked to provide spoken responses. Researchers initially administered the FS listening task, which he completed attentively and appeared to enjoy. However, when asked to retell the story, he was unable to generate an adequate verbal response. Based on this observation, researchers transitioned to a computer-based visual search task that emphasized visual attention, reaction time, and response accuracy. This maintained participant engagement and enabled successful completion of other tasks within our research protocol.

## 4. Discussion

### 4.1 Key Findings

In this retrospective study, we investigated the role of comorbidities in task-based, intracranial research participation and evaluated factors important in making the research experience inclusive and enjoyable for our patient participants. We found that broad inclusion is achievable, even when participants present with comorbidities and other barriers to participation. When conducting research involving patient participants, ethical integrity and practical sensitivity are crucial. In our experience with participants undergoing epilepsy surgery, we found that encouraging participant agency, asserting boundaries between clinical care and research, and reflecting and improving upon strategies for participant enjoyment, ensures a respectful and successful approach to research.

Motivators and deterrents to research participation have not been examined in pediatric epilepsy patients undergoing sEEG placement. However, related work in other underrepresented, medically complex populations provides useful context, as such groups face analogous recruitment and participation barriers [52–54], as does research in adult epilepsy patients undergoing intracranial monitoring or with implanted devices [45,47,55]. Zvolanek et al. [52] worked with patients with cerebral palsy, and found that schedule and travel limitations were the most prevalent barriers to participation in research, in addition to suggesting financial incentives to offset family burden. These barriers were similarly discussed in Renwick et al. [53], who point out that travel barriers can be even more burdensome to participants from rural communities. In our patient cohort, schedule and travel barriers were largely eliminated as recruitment occurred during in-patient hospitalization, though it is true that there would be selection bias in our cohort based on who is able to have brain surgery. In addition, while we did not provide compensation nor direct benefit to study participants, participants and caregivers reported primarily participating out of a sense of altruism. In a scoping review by Shariq et al. [54], the authors emphasized the need for person-centered approaches to consent that may help to overcome stereotypes that researchers may have regarding people with disabilities as poor candidates for research. They also describe a need for “co-design” of research protocols, in which community members and stakeholders are involved in designing and adapting protocols to participant needs. While our study did not directly utilize patient advocates in this way, by soliciting patient and family feedback on the research studies we were able to adapt protocols and understand their perspectives on motivations to participate. In work investigating ethics and barriers to participation for adult patients undergoing surgery for epilepsy, another consideration that we also emphasize here is the distinction between the research and clinical treatment [47,55,56]. When the clinical team is involved in research or consent, this can enhance trust, but can also blur boundaries between what is perceived as clinical treatment and basic science research. While the barriers to participation differ across these populations and settings, a common thread is the importance of flexible, responsive protocol design. This includes soliciting participant and family input and adapting procedures accordingly.

Our study shows that it is possible to design research studies that are inclusive of pediatric epilepsy patients with a broad spectrum of developmental delays. Passive tasks required no overt response and included listening to phonetically annotated movie trailers, narrated children’s stories, sentences, or syllables. Active tasks required spoken or behavioral responses, such as repeating consonant-vowel syllables or sentences embedded in noise, reading aloud, retelling a story from memory, or detecting target sounds in an attention task. Comorbidities did not appear to affect overall time spent doing research tasks and, aside from global developmental delay, did not affect participation in active tasks. This suggests patients need not be excluded based on these comorbidities, though such conditions may need to be incorporated into interpretation of results.

### 4.2 Suggestions for Task Design Improvements

To ensure research tasks can be completed by participants of varying cognitive and language abilities, we suggest several strategies for other researchers. The first is to incorporate natural, narrative stimuli rather than sentence lists, single word naming, or syllable listening, tasks that are typically less engaging for children [57]. By using modern tools to perform automatic annotation and forced alignment of phoneme and word boundaries, it is possible to replace typical linguistic corpora, as long as the stimuli are appropriately validated [58,59]. Second, multiple versions of a task could be provided: for example, production tasks that can accommodate different reading levels, as used by our group with these patients in prior work [41]. Third, researchers could consider tasks that may be used in multiple ways, such as tasks that include both production and perception components that can be used either separately or together. Fourth, researchers should design tasks that can be completed in short 5-10 minute blocks, with breaks in between. The nature of intracranial research is that interruptions can frequently occur, so building breaks into task design helps not only with finishing tasks, but allowing for inevitable disruptions. Finally, researchers should consider ways to gamify tasks to improve engagement [60].

### 4.3 Future Directions

Despite encouraging findings, there are still opportunities for improvement across the research engagement process. One area that might differ across research groups is the timing of consent. Our group has found that approaching the patients and their families two to three days after surgery has worked well and allowed families to make informed decisions considering their current clinical circumstances. Another approach could be to introduce the research during preoperative clinical consultations. This option should be considered if there are stipulations for the implantation of additional electrodes meant solely for research purposes, or if there are multiple research teams seeking to enroll the same participant.

Future task development may benefit from more explicitly incorporating feedback from participants and their caregivers. Our group currently administers a post research survey that includes basic questions regarding the participant’s experience in the study, including what they enjoyed or disliked. Adding survey questions about specific types of activities or media the participants are already engaging with outside of a hospital setting may allow researchers to model tasks after familiar and motivating formats. Still, careful consideration must be given to the need for stimulus consistency across participants, as too much variability in task design may limit comparability of results over time.

We hope that this work will serve as the foundation for the continued expansion of research with children undergoing Phase 2 surgical epilepsy monitoring with intracranial sEEG or ECoG. The strategies, perspectives, and considerations discussed herein should guide the consent, enrollment, and engagement of pediatric populations in research initiatives that are crucial to deepening our knowledge of the developing brain.

## Acknowledgments

The authors would like to thank Dr. Saskia Hendriks for helpful discussion and feedback on the manuscript. The authors would like to thank Maansi Desai, G. Lynn Kurteff, Elise Rickert, Rajvi Agravat, and Gerick Lee for assistance with task design, and Sadrita Mondal for assistance with compiling information for the case studies.

