## Supplementary Materials for "Perspectives in conducting task-based research in pediatric surgical epilepsy patients"

**Appendix A.** List of passive and active tasks used in our studies. Some of these have been used in previously published work [1–4], some are being incorporated into work in progress.

1. **Passive Tasks**
   a. *Movie Trailers:* Participants viewed annotated movie trailers from Pixar, Disney, and other studios. Trailers were phonetically transcribed and validated in prior scalp EEG work [1]. These stimuli were chosen to be engaging and enjoyable for a wide range of ages, while not feeling like a typical neuroscience task [1,5].
   b. *Sentence listening - TIMIT/Frog Story*: We incorporated two options for this task. (1) Participants listened to phonetically transcribed sentences from the TIMIT acoustic-phonetic corpus [6]. (2) As an alternative, we provided a narrated children’s story (“Frog, where are you?”) via iPad, offering participants a more engaging listening option. This story is part of an elicitation paradigm developed by the Systematic Analysis of Language Transcripts (SALT) software team (<https://www.saltsoftware.com/resources/elicaids/frogstories>) that has been used along with other “Frog stories” to study children’s language development [7].
2. **Active Tasks**
   a. *CV*: Participants listen to isolated consonant–vowel (CV) syllables and are asked to repeat them aloud.
   b. *Bamford-Kowal-Bench Speech-in-Noise (BKB-SIN)*: Participants listen to sentences presented with progressively increasing levels of 4-talker babble noise [8]. These stimuli are taken from a clinical audiology assessment for understanding speech in noise (Interacoustics, Inc.). After each sentence, they are asked to repeat it back as accurately as possible.
   c. *Sentence reading*: Read sentences aloud from the MOCHA-TIMIT database [9] or sentences modified from this database to be appropriate for a 3^rd^ grade reading level [4]. After each sentence is read aloud, the participant will hear playback of that sentence or a playback of a previously uttered sentence. Participants were excluded from this task if reading ability or parental input suggested it was not feasible, or if the participant attempted but did not wish to continue.
   d. *Delayed auditory feedback (DAF)*: Participants read aloud sentences and then listened to playback of their own voice.
   e. *Speech Motor Control*: Participants replicated specific facial motor movements taken from the American Speech-Language-Hearing Association Oral Mechanism Exam (<https://www.asha.org/practice-portal/clinical-topics/articulation-and-phonology/?srsltid=AfmBOooli3qtuJcMsc8tiwDduLaYewqE8QWsWEcoxdlZB3akqG4OTPf8#collapse_5>).
   f. *Visual Search*: Participants completed a timed computer-based visual discrimination task in which they identified a visual object that differed from the others, with both accuracy and reaction time recorded.
   g. *Frog Story Repeat*: Participants listened to the “Frog Story” (as in the passive task above) and were then instructed to retell the story aloud from memory, using page cues as recall aids.
   h. *Naturalistic Sounds*: Participants were presented with a continuous sequence of natural sounds from [10] and performed a 1-back task in which they were instructed to press a button whenever they detected a sound that had been repeated from the previous trial.
   i. *Auditory Attention*: Participants listened to a series of short movie trailer audio clips mixed with background music at varying signal-to-noise ratios. While listening, they were asked selectively attend to a specific target sound or word. Each time the target sound or word occurs during the clip, participants were asked to press a button as quickly and accurately as possible.
